# Rethinking dopamine as generalized prediction error

**DOI:** 10.1101/239731

**Authors:** Matthew P.H. Gardner, Geoffrey Schoenbaum, Samuel J. Gershman

## Abstract

Midbrain dopamine neurons are commonly thought to report a reward prediction error, as hypothesized by reinforcement learning theory. While this theory has been highly successful, several lines of evidence suggest that dopamine activity also encodes sensory prediction errors unrelated to reward. Here we develop a new theory of dopamine function that embraces a broader conceptualization of prediction errors. By signaling errors in both sensory and reward predictions, dopamine supports a form of reinforcement learning that lies between model-based and model-free algorithms. This account remains consistent with current canon regarding the correspondence between dopamine transients and reward prediction errors, while also accounting for new data suggesting a role for these signals in phenomena such as sensory preconditioning and identity unblocking, which ostensibly draw upon knowledge beyond reward predictions.

## Introduction

The hypothesis that midbrain dopamine neurons report a reward prediction error (RPE, the discrepancy between observed and expected reward) enjoys a seemingly unassailable accumulation of support from electrophysiology [1, 2, 3, 4, 5], calcium imaging [6, 7], optogenetics [8, 9, 10], voltammetry [11, 12], and human brain imaging [13, 14]. The success of the RPE hypothesis is exciting because the RPE is precisely the signal a reinforcement learning (RL) system would need to update reward expectations [15, 16]. Support for this RL interpretation of dopamine comes from findings that dopamine complies with basic postulates of RL theory [1], shapes the activity of downstream reward-predictive neurons in the striatum [17, 11], and plays a causal role in the control of learning [8, 9, 10, 13].

Despite these successes, however, there are a number of signs that this is not the whole story. First, it has long been known that dopamine neurons respond to novel or unexpected stimuli, even in the absence of changes in value [18, 19, 20, 7]. While some theorists have tried to reconcile this observation with the RPE hypothesis by positing that value is affected by novelty [21] or uncertainty [22], others have argued that this response constitutes a distinct function of dopamine [23, 24, 25], possibly mediated by an anatomically segregated projection from midbrain to striatum [7]. A second challenge is that some dopamine neurons respond to aversive stimuli. If dopamine responses reflect RPEs, then one would expect aversive stimuli to *reduce* responses (as observed in some studies; [26, 27]). A third challenge is that dopamine activity is sensitive to movement-related variables such as action initiation and termination [28, 29]. A fourth challenge is that dopamine activity [30] and its putative hemodynamic correlates [31] are influenced by information, such as changes in stimulus contingencies, that should in principle be invisible to a pure “model-free” RL system that updates reward expectations using RPEs. This has led to elaborations of the RPE hypothesis according to which dopamine has access to some “model-based” information, for examples in terms of probabilistic beliefs or samples from a model-based simulator [32, 33, 22, 34, 35, 36].

While some of these puzzles can be resolved within the RPE framework by modifying assumptions about the inputs to and modulators of the RPE signal, recent findings have proven more unyielding. In this paper we focus on three of these findings: (1) dopamine transients are necessary for learning induced by unexpected changes in the sensory features of expected rewards [37]; (2) dopamine neurons respond to unexpected changes in sensory features of expected rewards [38]; and (3) dopamine transients are both sufficient and necessary for learning stimulus-stimulus associations [39]. Taken together, these findings seem to contradict the RPE framework supported by so much other data.

Here we will suggest one possible way to reconcile the new and old findings, based on the idea that dopamine computes prediction errors over sensory features, much as was previously hypothesized for rewards. This sensory prediction error (SPE) hypothesis is motivated by normative considerations: SPEs can be used to estimate a predictive feature map known as the *successor representation* (SR; [40, 41]). The key advantage of the SR is that it simplifies the computation of future rewards, combining the efficiency of model-free RL with some of the flexibility of model-based RL. Neural and behavioral evidence suggests that the SR is part of the brain’s computational repertoire [42, 43], possibly subserved by the hippocampus [44, 45]. Here, building on the pioneering work of Suri [46], we argue that dopamine transients previously understood to signal RPEs may instead constitute the SPE signal used to update the SR.

## Theoretical framework

### The reinforcement learning problem

RL theories posit an environment in which an animal accumulates rewards as it traverses a sequence of “states” governed by a transition function *T*(*s*’|*s*), the probability of moving from state *s* to state *s*’, and a reward function *R*(*s*), the expected reward in state *s*. The RL problem is to predict and optimize *value*, defined as the expected discounted future return (cumulative reward):

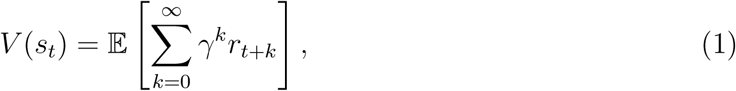

where *r*_*t*_ is the reward received at time *t* in state *s*_*t*_, and γ ∈ [0, 1] is a discount factor that determines the weight of temporally distal rewards. Because the environment is assumed to obey the Markov property (transitions and rewards depend only on the current state), the value function can be written in a recursive form known as the *Bellman equation* [47]:

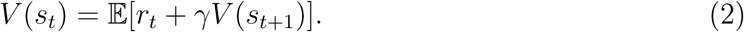

The Bellman equation allows us to define efficient RL algorithms for estimating values, as we explain next.

### Model-free and model-based learning

Model-free algorithms solve the RL problem by directly estimating *V* from interactions with the environment. The Bellman equation specifies a recursive consistency condition that the value estimate 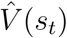 must satisfy in order to be accurate. By taking the difference between the two sides of the Bellman equation, 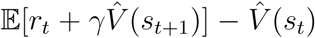, we can obtain a measure of expected error; the direction and degree of the error is informative about how to correct 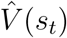.

Because model-free algorithms do not have access to the underlying environment model (*R* and *T*) necessary to compute the expected error analytically, they typically rely on a stochastic sample of the error based on experienced transitions and rewards:

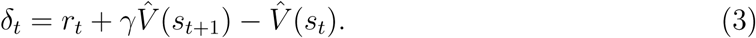

This quantity, commonly known as the *temporal difference (TD) error*, will on average be 0 when the value function has been perfectly estimated. The TD error is the basis of the classic TD learning algorithm [47], which in its simplest form updates the value estimate according to 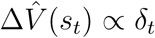. The RPE hypothesis states that dopamine reports the TD error [15, 16].

Model-free algorithms like TD learning are efficient because they *cache* value estimates, which means that state evaluation (and by extension action selection) can be accomplished by simply inspecting the values cached in the relevant states. This efficiency comes at the cost of flexibility: if the reward function changes at a particular state, the entire value function must be re-estimated, since the Bellman equation implies a coupling of values between different states. For this reason, it has been proposed that the brain also makes use of model-based algorithms [48, 49], which occupy the opposite end of the efficiency-flexibility spectrum. Model-based algorithms learn a model of the environment (*R* and *T*) and use this model to evaluate states, typically through some form of forward simulation or dynamic programming. This approach is flexible, because local changes in the reward or transition functions will instantly propagate across the entire value function, but at the cost of relying on comparatively inefficient simulation.

Some of the phenomena that we discuss in the Results have been ascribed to model-based computations supported by dopamine [50], thus transgressing the clean boundary between the model-free function of dopamine and putatively non-dopaminergic model-based computations. The problem with this reformulation is that it is unclear what exactly dopamine is contributing to model-based learning. Although prediction errors are useful for updating estimates of the reward and transition functions used in model-based algorithms, these do not require a TD error. A distinctive feature of the TD error is that it bootstraps a future value estimate (the 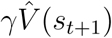 term); this is necessary because of the Bellman recursion. But learning reward and transition functions in model-based algorithms can avoid bootstrapping estimates because the updates are local thanks to the Markov property.

To make this concrete, a simple learning algorithm (guaranteed to converge to the maximum likelihood solution under some assumptions about the learning rate) is to update the model parameters according to:

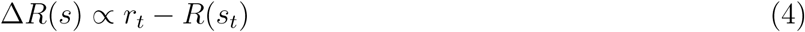

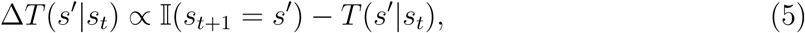

where 𝕀(·) = 1 if its argument is true, and 0 otherwise [51]. These updates can be understood in terms of prediction errors, but *not* TD errors (they do not bootstrap future value estimates). The TD interpretation is important for explaining phenomena like the shift in signaling to earlier reward-predicting cues [16], the temporal specificity of dopamine responses [52, 53], and the sensitivity to long-term values [54]. Thus, it remains mysterious how to retain the TD error interpretation of dopamine, which has been highly successful as an empirical hypothesis, while simultaneously accounting for the sensitivity of dopamine to SPEs.

### The successor representation

To reconcile these data, we will develop the argument that dopamine reflects sensory TD errors, encompassing both reward and non-reward features of a stimulus. In order to introduce some context to this idea, let us revisit the fundamental efficiency-flexibility trade-off. One way to find a middle-ground between the extremes occupied by model-free and model-based algorithms is to think about different ways to *compile* a model of the environment. By analogy with programming, a compiled program gains efficiency (in terms of runtime) at the expense of flexibility (the internal structure of the program is no longer directly accessible). Model-based algorithms are maximally uncompiled: they explicitly represent the parameters of the model, thus providing a representation that can be flexibly altered for new tasks. Model-free algorithms are maximally compiled: they only represent the summary statistics (state values) that are needed for reward prediction, bypassing a flexible representation of the environment in favor of computational efficiency.

A third possibility is a partially compiled model. [40] presented one such scheme, based on the following mathematical identity:

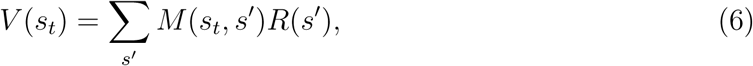

where *M* denotes the successor representation (SR), the expected discounted future state occupancy:

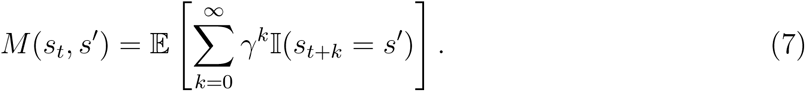

Intuitively, the SR represents states in terms of the frequency of their successor states. From a computational perspective, the SR is appealing for two reasons. First, it renders value computation a linear operation, yielding efficiency comparable to model-free evaluation. Second, it retains some of the flexibility of model-based evaluation. Specifically, changes in rewards will instantly affect values because the reward function is represented separately from the SR. On the other hand, the SR will be relatively insensitive to changes in transition structure, because it does not explicitly represent transitions—these have been compiled into a convenient but inflexible format. Behavior reliant upon such a partially-compiled model of the environment should be more sensitive to reward changes than transition changes, a prediction recently confirmed in humans [42].

The SR obeys a recursion analogous to the Bellman equation:

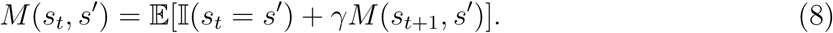

Following the logic of the previous section, this implies that a TD learning algorithm can be used to estimate the SR:

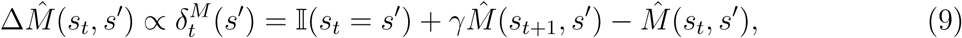

where 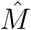 denotes the approximation of *M*.

One challenge facing this formulation is the *curse of dimensionality*: in large state spaces it is impossible to accurately estimate the SR for all states. Generalization across states can be achieved by defining the SR over state features (indexed by *j*) and modeling this feature-based SR with linear function approximation:

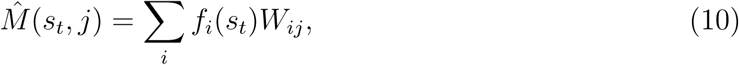

where *ƒ*_*i*_(*s*) denotes the *i*th feature of state *s* and *W* is a weight matrix that parametrizes the approximation. In general the features can be arbitrary, but for the purposes of this paper, we will assume that the features correspond to distinct stimulus identities; thus *ƒ*_*i*_(*s*) = 1 if stimulus *i* is present in state *s*, and 0 otherwise. Linear function approximation leads to the following learning rule for the weights:

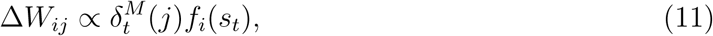

where

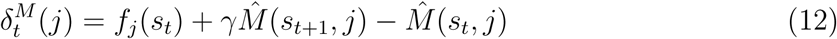

is the TD error under linear function approximation. We will argue that dopamine encodes this TD error.

One issue with comparing this vector-valued TD error to experimental data is that we don’t yet know how particular dopamine neurons map onto particular features. In order to make minimal assumptions, we will assume that each neuron has a uniform prior probability of encoding any given feature. Under ignorance about feature tuning, the expected TD error is then proportional to the superposition of feature-specific TD errors, 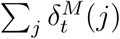. In our simulations of dopamine, we take this superposition to be the “dopamine signal” (see also [32]), but we wish to make clear that this is a provisional assumption that we ultimately hope to replace once the feature tuning of dopamine neurons is better understood.

There are several notable aspects of this new model of dopamine. First, it naturally captures SPEs, as we will illustrate shortly. Second, it also captures RPEs if reward is one of the features. Specifically, if *ƒ*_*j*_(*s*_*t*_) = *r*_*t*_, then the correspond column of the SR is equivalent to the value function, *M*(*s*, *j*) = *V* (*s*), and the corresponding TD error is the classical RPE, 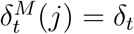. Third, the TD error is now vector-valued, which means that dopamine neurons may be heterogeneously tuned to particular features (as hypothesized by some authors; [55]), or they multiplex several features [56], or both. Notably, although the RPE correlate has famously been evident in single-units, representation of these more complex or subtle prediction errors may be an ensemble property.

## Simulations

Some of the most direct evidence for our hypothesis comes from a recent study by Chang et al. [37], who examined whether dopamine is necessary for learning about changes in reward identity (Figure 1A). Animals first learned to associate two stimuli (X_*B*_ and X_*UB*_) with different reward flavors. These stimuli were then reinforced in compound with other stimuli (A_*B*_ and A_*UB*_). Critically, the X_*UB*_A_*UB*_ trials were accompanied by a change in reward flavor, a procedure known as “identity unblocking” that attenuates the blocking effect [57, 58, 59]. This effect eludes explanation in terms of model-free mechanisms, but is naturally accommodated by the SR since changes in reward identity induce sensory prediction errors. Chang et al. [37] showed that optogenetic inhibition of dopamine at the time of the flavor change prevents this unblocking effect (Figure 1B). Our model accounts for this finding (Figure 1C), because inhibition suppresses SPEs that are necessary for driving learning.

**Figure 1:**
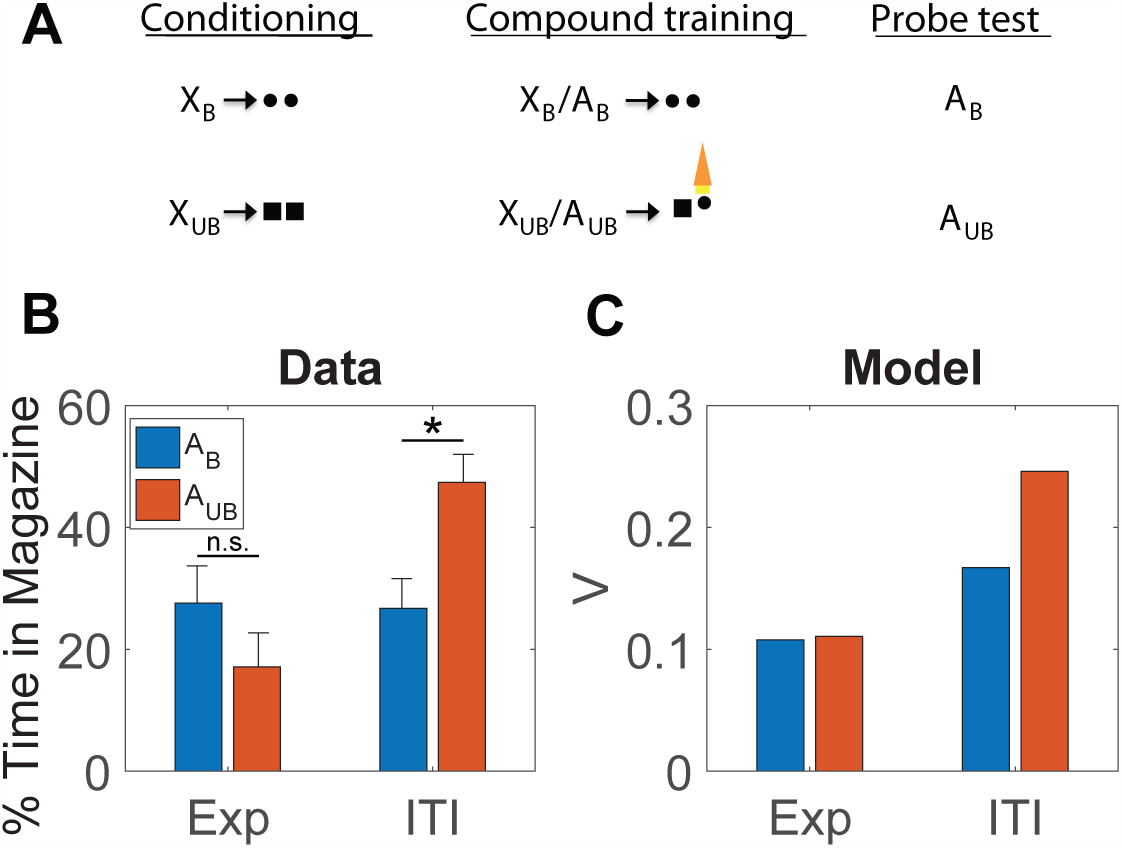
Inhibition of dopamine neurons prevents learning induced by changes in reward identity. (*A*) Identity unblocking paradigm. Circles and squares denote distinct reward flavors. Orange light symbol indicates when dopamine neurons were suppressed optogenetically to disrupt any positive SPE; this spanned a 5s period beginning 500ms prior to delivery of the second reward. (*B*) Conditioned responding on the probe test. Exp: experimental group, receiving inhibition during reward outcome. ITI: control group, receiving inhibition during the intertrial interval. Asterisk indicates significant difference (*p* < 0.05). Error bars show standard error of the mean. Data replotted from [37]. (*C*) Model simulation of the value function.

One discrepant observation is a simulated increase in V in the ITI condition relative to the Exp condition for A_*B*_, which does not appear in the experimental data. During the second stage of learning, X_*UB*_, A_*UB*_, and the sensory features of both pellet types are presented together. Because of the co-occurrence of these features, associations develop between them such that the sensory features of the pellets now have slight associations with one another as well as the cues that predict them. This causes A_*B*_ and X_*B*_ to have a slight association with the sensory features of the pellet that it never predicted since both pellets now have mild associations with one another.

Electrophysiological experiments have confirmed that dopamine neurons respond to changes in identity, demonstrating a neural signal that is capable of explaining the data from Chang et al. [37]. We have already mentioned the sizable literature on novelty responses, but the significance of this activity is open to question, because the animal’s prior value expectation is typically unclear. A study reported by Takahashi et al. [38] provides more direct evidence for an SPE signal, using a task (Figure 2A) in which animals experience both shifts in value (amount of reward) and identity (reward flavor). Takahashi and colleagues found that individual dopamine neurons exhibited the expected changes in firing to shifts in value (Figure 2B, reward addition and omission) and also showed a stronger response following a value-neutral change in reward identity (Figure 2B, identity switch), changes in firing similar to those predicted by the model under these conditions (Figure 2C).

**Figure 2:**
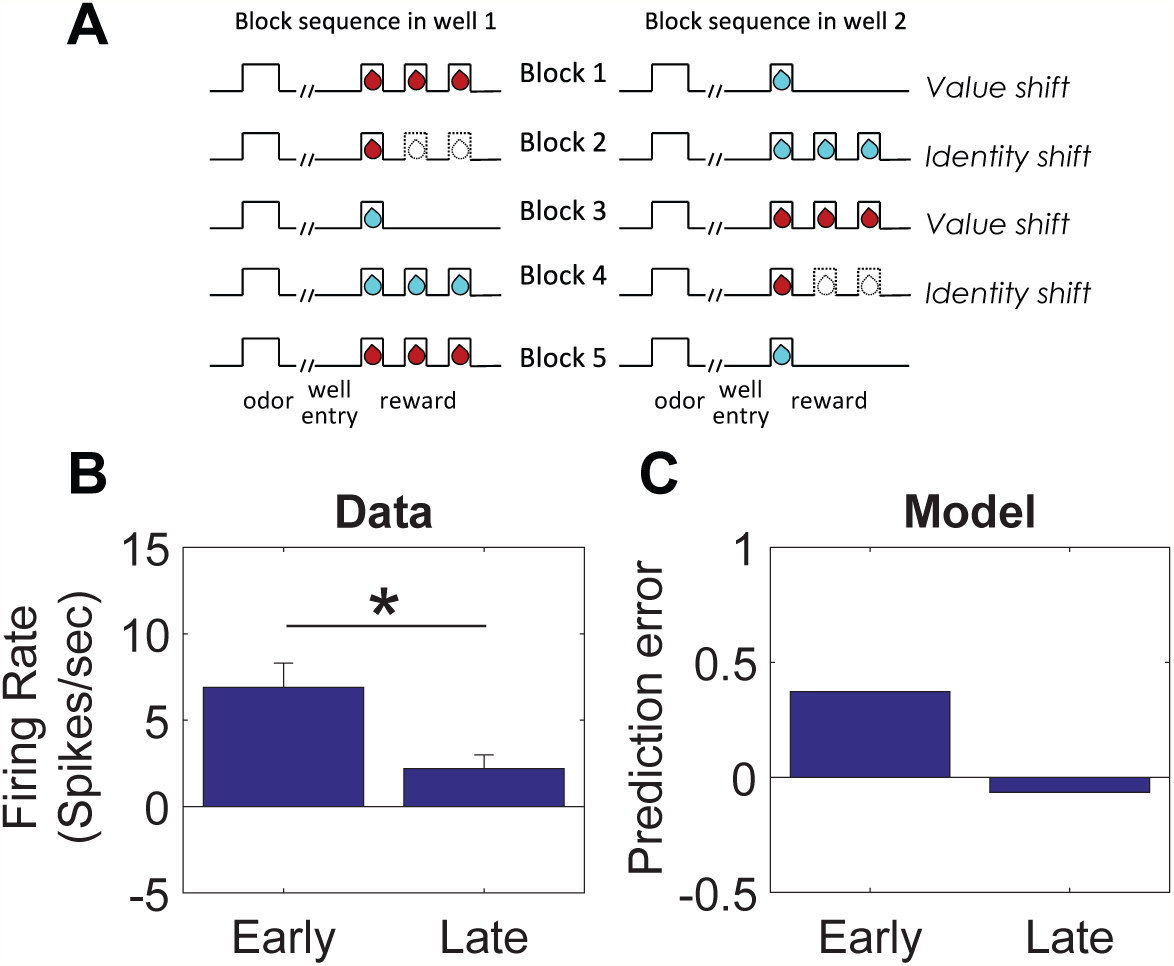
Dopamine neurons respond to changes in reward identity. (*A*) Time course of stimuli presented to the animal on each trial. Dashed indicate reward omission, solid lines indicate reward delivery. At the start of each session, one well was randomly designated as short (a .5-s delay before the reward) and the other, long (a 1- to 7-s delay before the reward; see Block 1). In Block 2, these contingencies were switched. In Block 3, the delay was held constant, while the number of rewards was manipulated; one well was designated a big reward, in which a second bolus of reward was delivered (big reward), and a small (single bolus) reward was delivered in the other well. In Block 4, these contingencies were switched again.(*B*) Firing rate of dopamine neurons on trials that occurred early (first 5 trials) or late (last 5 trials) during an identity shift block. Error bars show standard error of the mean. Data replotted from [38]. (*C*) Model simulation of TD error.

A strong form of our proposal is that dopamine transients are both sufficient and necessary for learning stimulus-stimulus associations. Recent experiments using a sensory preconditioning paradigm [39] have tested this using sensory preconditioning. In this paradigm (Figure 3A), various stimuli and stimulus compounds (denoted A, EF, AD, AC) are associated with another stimulus X through repeated pairing in an initial preconditioning phase. In a subsequent conditioning phase, X is associated with reward (sucrose pellets). In a final probe test, conditioned responding to a subset of the individual stimuli (F, D, C) is measured in terms of the number of food cup entries elicited by the presentation of the stimuli. During the preconditioning phase, one group of animals received optogenetic activation of dopamine neurons via channelrhodopsin (ChR2) expressed in the ventral tegmental area of the midbrain. In particular, optogenetic activation was applied either coincident with the onset of X on AC→X trials, or (as a temporal control) 120-180 seconds after X on AD→X trials. Another control group of animals received the same training and optogenetic activation, but expressed light-insensitive enhanced yellow fluorescent protein (eYFP).

**Figure 3:**
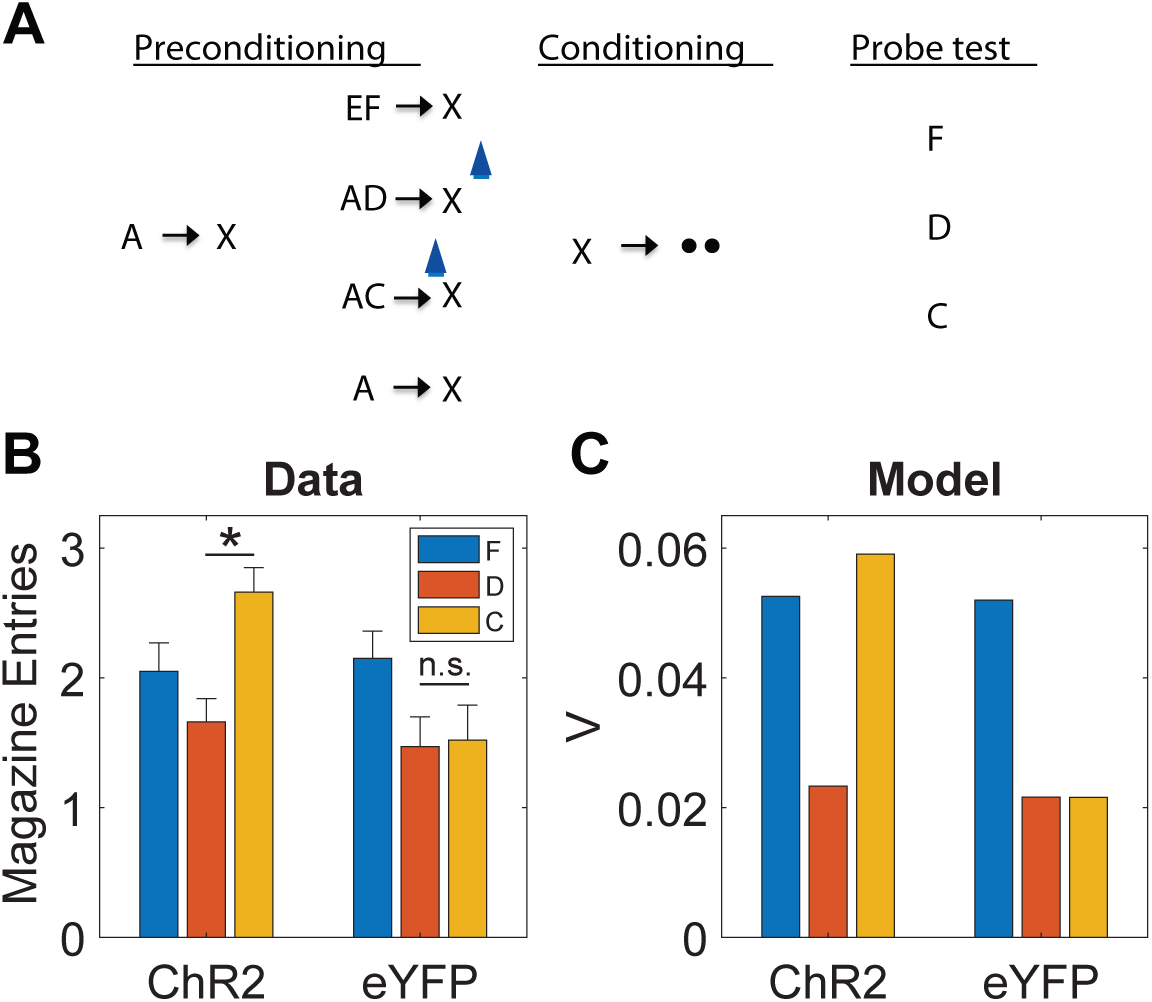
Dopamine transients are sufficient for learning stimulus-stimulus associations. (*A*) Sensory preconditioning paradigm. The initial preconditioning phase is broken down into two sub-phases. Letters denote stimuli, arrows denote temporal contingencies, and circles denote rewards. Blue light symbol indicates when dopamine neurons were activated optogenetically to mimic a positive SPE; this spanned a 1s period beginning at the start of X. (*B*) Number of food cup entries occurring during the probe test for experimental (ChR2) and control (eYFP) groups. Error bars show standard error of the mean. Data re-plotted from [39]. (*C*) Model simulation, using the value estimate as a proxy for conditioned responding.

A blocking effect was discernible in the control (eYFP) group, whereby A reduced acquisition of conditioned responding to C and D, compared to F, which was trained in compound with a novel stimulus (Figure 3B). The blocking effect was eliminated by optogenetic activation in the experimental (ChR2) group, specifically for C, which received activation coincident with X. Thus, activation of dopamine neurons was sufficient to drive stimulus-stimulus learning in a temporally specific manner.

These findings raise a number of questions. First, how does one explain blocking of stimulus-stimulus associations? Second, how does one explain why dopamine affects this learning in the apparent absence of new reward information?

In answer to the first question, we can appeal to an analogy with blocking of stimulus-reward associations. The classic approach to modeling this phenomenon is to assume that each stimulus acquires an independent association and that these associations summate when the stimuli are presented in compound [60]. While there are boundary conditions on this assumption [61], it has proven remarkably successful at capturing a broad range of learning phenomenon, and is inherited by TD models with linear function approximation (e.g., [16, 22, 62]). Summation implies that if one stimulus (A) perfectly predicts reward, then a second stimulus (C) with no pre-existing association will fail to acquire an association when presented in compound with A, because the sum of the two associations will perfectly predict reward and hence generate an RPE of 0. The same logic can be applied to stimulus-stimulus learning by using linear function approximation of the successor representation, which implies that stimulus-stimulus associations will summate and hence produce blocking, as observed in Sharpe et al. [39].

In answer to the second question, we argue that dopamine is involved in stimulus-stimulus learning because it reflects a multifaceted SPE, as described in the previous section. By assuming that optogenetic activation adds a constant to the SPE (see Methods), we can capture the unblocking findings reported by Sharpe and colleagues (Figure 3C). The mechanism by which optogenetic activation induces unblocking is essentially the same as the one suggested by the results of Steinberg et al. [9] for conventional stimulus-reward blocking: by elevating the prediction error, a learning signal is engendered where none would exist otherwise. However, while the results of Steinberg and colleagues are consistent with the original RPE hypothesis of dopamine, the results of Sharpe et al. [39] cannot be explained by this model and instead require the analogous dopamine-mediated mechanism for driving learning with SPEs.

In addition to establishing the sufficiency of dopamine transients for learning, [39] also established their necessity, using optogenetic inactivation. In a variation of the sensory preconditioning paradigm (Figure 4A), two pairs of stimulus-stimulus associations were learned (A→X and B→Y). Subsequently, X and Y were paired with different reward flavors, and finally conditioned responding to A and B was evaluated in a probe test. In one group of animals expressing halorhodopsin in dopamine neurons (NpHR), optogenetic inhibition was applied coincident with the transition between the stimuli on B→Y trials. A control group expressing light-insensitive eYFP was exposed to the same stimulation protocol. Sharpe and colleagues found that inhibition of dopamine selectively reduced responding to B (Figure 4B), consistent with our model prediction that disrupting dopamine transients (a negative prediction error signal) should attenuate stimulus-stimulus learning (Figure 4C).

**Figure 4:**
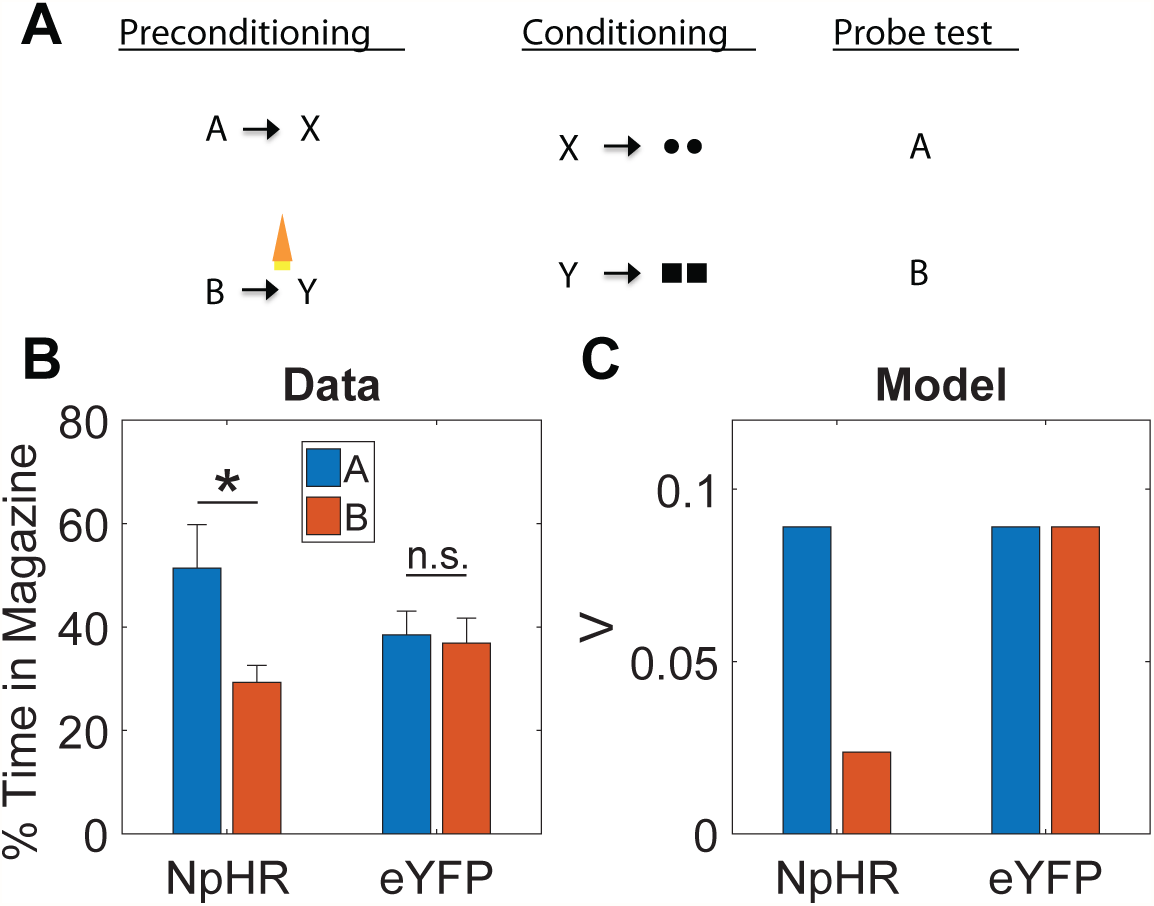
Dopamine transients are necessary for learning stimulus-stimulus associations. (*A*) Sensory preconditioning paradigm. Circles and squares denote distinct reward flavors. Orange light symbol indicates when dopamine neurons were suppressed optogeneti-cally to disrupt any positive SPE; this spanned a 2.5s period beginning 500ms prior to the end of B. (*B*) Number of food cup entries occurring during the probe test for experimental (NpHR) and control (eYFP) groups. Error bars show standard error of the mean. Data replotted from [39]. (*C*) Model simulation.

## Limitations and extensions

One way to drive a wedge between model-based and model-free algorithms is to devalue rewards (e.g., through pairing the reward with illness or selective satiation) and show effects on previously acquired conditioned responses to stimuli that predict those rewards. Because model-free algorithms like TD learning need to experience unbroken stimulus-reward sequences to update stimulus values, the behaviors they support are insensitive to such reward devaluation. Model-based algorithms, in contrast, are able to propagate the devaluation to the stimulus without direct experience, and hence allow behavior to be devaluation-sensitive. Because of this, devaluation-sensitivity has frequently been viewed as an assay of model-based RL [48].

However, such sensitivity can also be a property of SR-based RL, since the SR represents the association between the stimulus and food and is also able to update the reward function of the food as a result of devaluation. Thus, like model-based accounts, an SR model can account for changes in previously learned behavior to reward-predicting stimuli after devaluation, both in normal situations [43, 42] and when learning about those stimuli is unblocked by dopamine activation [63]. However, the SR model cannot spontaneously acquire transitions between states that are not directly experienced [43, 42]. With this in mind, we consider the finding that reward devaluation alters the learning induced by activation of dopamine neurons in the sensory preconditioning paradigm of Sharpe et al. [39].

A key aspect of the reward devaluation procedure is that the food was paired with illness after the end of the entire preconditioning procedure and in the absence of any of the stimuli (and in fact not in the training chamber). In the SR model, only stimuli already predictive of the food can change their values after devaluation. In the paradigm of Sharpe and colleagues, X was associated with food but C was not. Moreover, C was associated with X before any association with food was established. Because of this, C is not updated in the SR model to incorporate an association with food. It follows that, unlike the animals in Sharpe et al. [39], the model will not be devaluation-sensitive when probed with C (Figure 5B).

**Figure 5:**
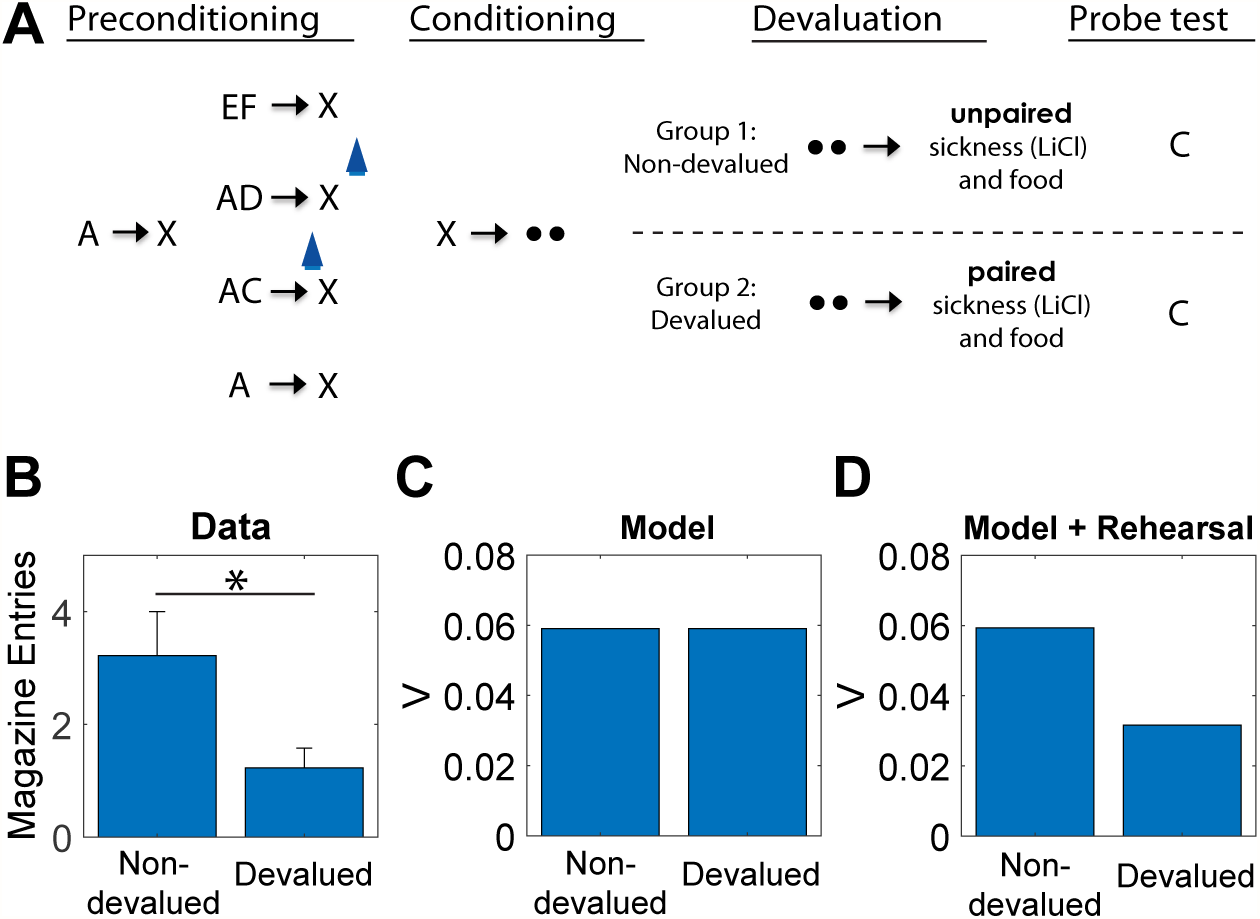
Behavior to preconditioned cue that is unblocked by activation of dopamine neurons is sensitive to devaluation of the predicted reward. Data (A, replotted from [39]) and model simulation (B) for conditioned responding to stimulus C in the probe test. Animals in the devalued group were injected with lithium chloride in conjunction with ingestion of the reward (sucrose pellets), causing a strong aversion to the reward. Animals in the nondevalued group were injected with lithium chloride approximately 6 hours after ingestion of the reward. Error bars show standard error of the mean. (C) A version of the model with rehearsal of stimulus X during reward devaluation was able to capture the devaluation-sensitivity of animals.

It is possible to address this failure within our theoretical framework in a number of different ways. One way we considered was to allow optogenetic activation to increment predictions for *all* possible features, instead of being restricted to recently active features by a *feature eligibility trace* (see Methods), as in the simulations thus far. With such a promiscuous artificial error signal, the model can recapitulate the devaluation effect, because C would then become associated with food (along with everything else) in the preconditioning phase itself. The problem with this work-around is that it also predicts that animals should develop a conditioned response to the food cup for all the cues during preconditioning, since food cup shaping prior to preconditioning seeds the food state with reward value. As a result, any cue paired with the food state immediately begins to induce responding at the food cup. Such behavior is not observed, suggesting that the artificial update caused by optogenetic activation of the dopamine neurons is locally restricted.

A second more conventional way to address this failure within our theoretical framework is to assume that there is some form of offine rehearsal or simulation that is used to update cached predictions [33, 64, 65]. Russek et al. [43] have shown that such a mechanism is able to endow SR-based learning with the ability to retrospectively update predictions even in the absence of direct experience. A minimal implementation of such a mechanism in our model, simply by “confabulating” the presence of X during reward devaluation, is sufficient to capture the effects of devaluation following optogenetic activation of dopamine neurons (Figure 5C). This solution makes the experimental prediction that the devaluation-sensitivity of this artificially unblocked cue should be time-dependent, under the assumption that the amount of offine rehearsal is proportional to the retention interval.

## Discussion

The RPE hypothesis of dopamine has been one of theoretical neuroscience’s signature success stories. This paper has set forth a significant generalization of the RPE hypothesis that enables it to account for a number of anomalous phenomena, without discarding the core ideas that motivated the original hypothesis. The proposal that dopamine reports a SPE is grounded in a normative theory of reinforcement learning [40], motivated independently by a number of computational [43, 66], behavioral [42, 67, 68] and neural [44, 45, 69, 70] considerations.

An important strength of the proposal is that it extends the functional role of dopamine beyond RPEs, while still accounting for the data that motivated the original RPE hypothesis. This is because, if reward is treated as a sensory feature, then one dimension of the vector-valued SPE will be the RPE. Indeed, dopamine SPEs should behave systematically like RPEs, except that they respond to features: they should pause when expected features are unexpectedly omitted, they should shift back to the earliest feature-predicting cue, and they should exhibit signatures of cue competition, such as overexpectation. SPEs are used to update cached predictions, analogous to the RPE in model-free algorithms. However these cached predictions extend beyond value to include information about the occupancy of future states (the SR). The SR can be used in a semi-flexible manner that allows behavior to be sensitive to changes in the reward structure, such as devaluation by pairing a reward with illness. As a result, even if dopamine is constrained by the model proposed here, it would support significantly more flexible behavior than supposed by classical model-free accounts [15, 16], even without moving completely to an account of model-based computation in the dopamine system [50].

Nevertheless, the theory proposed here—particularly if it incorporates offline rehearsal in order to fully explain the results of Sharpe et al. [39]—does strain the dichotomy between model-based and model-free algorithms that has been at the heart of modern RL theories [48]. However, as noted earlier, SR requires offine rehearsal to incorporate the effects of devaluation after preconditioning in Sharpe et al or manipulations of the transition structures of tasks [42]. If these effects, and particularly dopamine’s involvement in them, are mediated by an SR mechanism, then we should be able to interfere with it by manipulating retention intervals or attention [33]. For example, the strength of devaluation sensitivity in Sharpe et al should be diminished by a very short retention interval prior to the probe test, since this would reduce time available for rehearsal. If these effects do not show any dependence on the length of the retention interval, then this would be more consistent with model-based algorithms, which do not require any rehearsal.

Another testable prediction of the theory is that we should see heterogeneity in the dopamine response, reflecting the vector-valued nature of the SPE. Importantly, such tuning need not be statistically evident in the spiking of an individual neuron. It might show up in the pattern of response across the entire population or even in subpopulations determined by target or other criteria. Indeed, target-based heterogeneity is already evident in some studies of dopamine release or function in downstream regions [63, 71, 72]. Related to this, the theory also predicts the existence of a negative SPE to allow reductions in the strength of weights in the SR. In its simplest form, the omission of an expected stimulus could result in suppression of firing, analogous to reward omission responses [16]. However, this effect might be subtle if SPEs are population-coded by the dopamine signal, as suggested above; the negative SPE may simply reflect a particular pattern across the population rather than overt suppression at the level of single neurons. Distinctive patterns of activity identifying the source of the error and differentiating the addition of information versus its omission sets our proposal apart from explanations based on salience signals, which are typically thought to be both non-specific and unsigned [73]. These predictions set an exciting new agenda for dopamine research by embracing a broader conception of dopamine function.

While we have focused on dopamine in this paper, a complete account will obviously need to integrate the computational functions of other brain regions. Where does information relevant to computing SPE’s come from? One obvious possibility is from sensory regions. Sensory areas respond both to and in expectation of external events [?, ?, ?], and these areas send input to brainstem, thus they are positioned to feed information to the dopaminergic system. Beyond this, the hippocampus and orbitofrontal cortex seem likely to be particularly important. Many lines of evidence are consistent with the idea that the hippocampus encodes a “predictive map” resembling the SR [44]. For example, hippocampal place cells alter their tuning with repeated experience to fire in anticipation of future locations [74], and fMRI studies have found predictive coding of non-spatial states [45, 75]. The orbitofrontal cortex has also been repeatedly implicated in predictive coding, particularly of reward outcomes [76, 77], but also of sensory events [78, ?], and the orbitofrontal cortex is critical for sensory-specific outcome expectations in Pavlovian conditioning [79]. Wilson et al. [80] have proposed that the orbitofrontal cortex encodes a “cognitive map” of state space, which presumably underpins this diversity of stimulus expectations. Thus, evidence suggests that both hippocampus and orbitofrontal cortex encode some form of predictive representation [81]. Further, dopaminergic modulation of these regions is well-established [82, 83]. It is tempting to speculate that afferent input from and dopaminergic modulation of the hippocampus and orbitofrontal cortex may be especially critical to the SPE function proposed here.

The influence of these representations may be filtered through interactions with more basic value representations in striatum. This proposal fits with the observation that the hippocampus and orbitofrontal cortex appear to confer stimulus specificity on value-sensitive neurons in the striatum [84, 85]. Striatal value representations are already proposed to influence activity in VTA [86]. By this model, dopamine would still provide the RPE signal that drives striatal plasticity related to actions or “value”, as in most contemporary accounts, but in addition it would provide an SPE signal to update associative representations, perhaps in striatum but also in upstream orbitofrontal and hippocampal areas, which feed into the striatum. While speculative, this idea is consistent with findings showing heterogeneity of dopamine function based on projection target, at least within striatum [63, 71]. It is also consistent with recent human imaging work, confirming the presence of an SPE-like signal in human VTA, and reporting that the strength of this signal during learning is correlated with the strength of new sensory-sensory correlates developed in the orbitofrontal cortex [70].

The view that dopamine reports the SR prediction error provides a bridge between sensory and reward prediction error accounts of dopamine function. The tension between these views has long vexed computational theories, and has posed particular problems for pure RPE accounts of dopamine. We see our model as the first step towards resolving this tension. While we have shown that the notion of a generalized prediction error is consistent with a wealth of empirical data, this is just the beginning of the empirical enterprise. Armed with a quantitative framework, we can now pursue evidence for such prediction errors with greater precision and clarity.

## Methods

### Linear value function approximation

Under the linear function approximation scheme described in the Results, the value function estimate is given by:

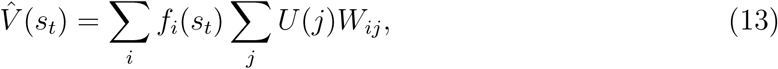

where *U*(*j*) is the reward expectation for feature *j*, updated according to a delta rule:

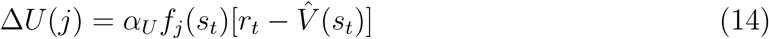

with learning rate *α*_*U*_.

In the supplementary figures, we report simulations of the value-based TD learning algorithm, TD(0), which approximates the value function using linear function approximation:

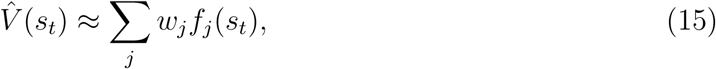

and updates the weights according to:

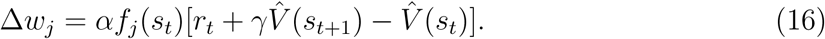

### Excitatory and inhibitory asymmetry in the TD error term

There is a large body of evidence in associative learning suggesting an imbalance between excitatory and inhibitory learning [87, 88]. Mirroring this imbalance is an asymmetry in the dynamic range of the firing rate of single dopaminergic neurons in the midbrain [2]. In accordance with these observations, we assume that the error terms (Δ*W*_*ij*_ and Δ*U*_*j*_) are rescaled by a factor of 1/4 for negative prediction errors. This is equivalent to assuming separate learning rates for positive and negative prediction errors [89]. Note that, following prior theoretical work (e.g., [16]), we consider negative prediction errors to be coded by real neurons relative to a baseline firing rate, acknowledging the fact that neurons cannot produce negative firing rates.

### Simulation parameters

We used the following parameters in the simulations of SR: γ = 0.95, *α*_*W*_ = 0.06, *α*_*U*_ = 0.03, where *α*_*W*_ is the learning rate for the weight matrix *W*, *α*_*U*_ is the learning rate for the reward function, and γ is the discount rate. For the model-free TD learning algorithm simulations, we used the following parameters: γ = 0.95, *α* = 0.05. We used the same set of parameters across all simulations. However, our results are largely robust to variations in these parameters.

### Modeling optogenetic activation and inhibition

Optogenetic intervention was modeled by modifying the TD error as follows:

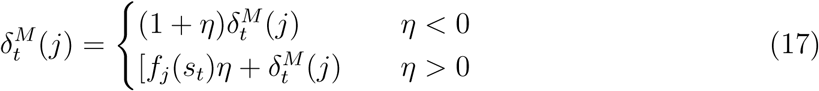

where *η* = 1.0 for optogenetic activation and –0.8 for inhibition. The asymmetry between the functions for activation and inactivation was chosen to better match the the hypothesized function of optogenetic stimulation based on empirical findings. For positive stimulation of dopamine, it is thought that the increased dopamine activity should enhance learning with the currently active features, which in the SR model is the *ƒ*_*j*_(*s*_*t*_) term. For optogenetic inhibition of dopamine, we have found that punctate versus prolonged inhibition causes differential effects, with punctate inhibition resulting in negative prediction errors and prolonged inhibition resulting in shunting of the error signal [10]. Our inhibition in the experiments included in this paper were prolonged, necessitating a different model of the inhibitory optogenetic manipulation.

## Acknowledgments

We are grateful to Yael Niv, Mingyu Song, Brian Sadacca and Andrew Wikenheiser for helpful discussions.

## Ethics statement

Not applicable.

## Data accessibility statement

All simulation code is available at https://github.com/mphgardner/TDSR.

## Authors’ contribution statement

MPHG, SJG and GS conceived the ideas and wrote the manuscript. MPHG and SJG carried out the model simulations and created the figures.

## supplementary figures

**Figure 1:**
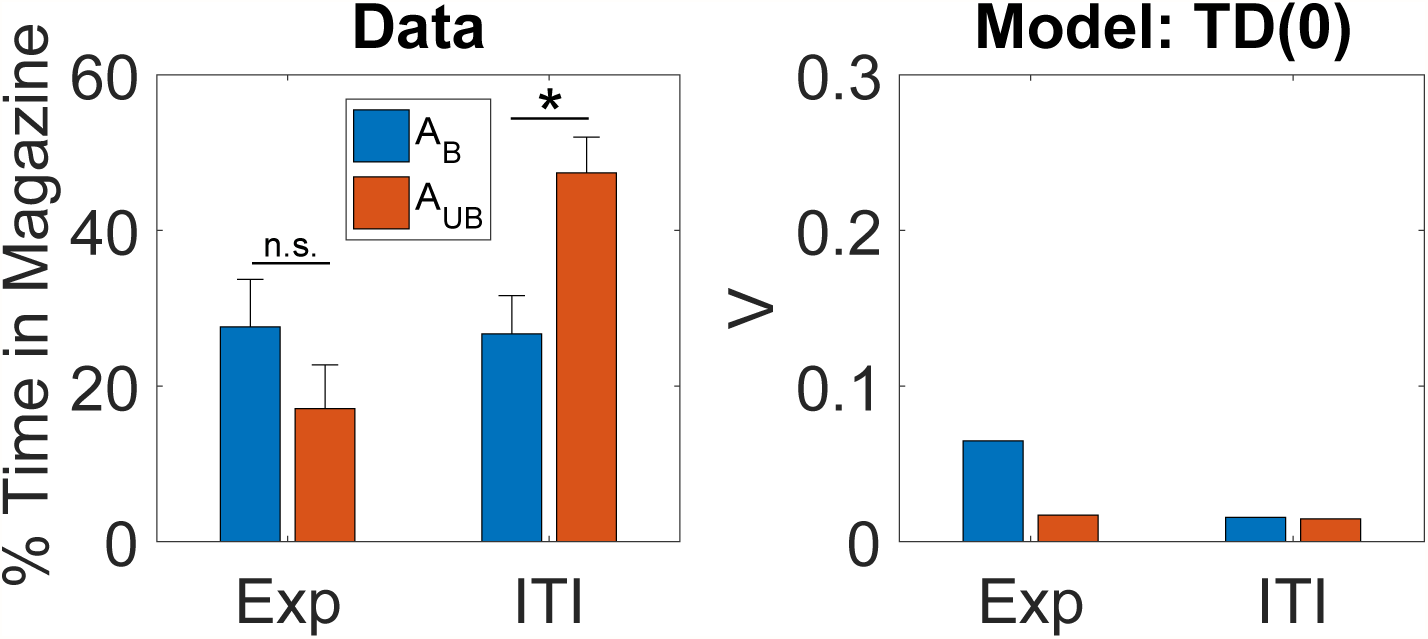
Inhibition of dopamine neurons prevents learning induced by changes in reward identity. (*Left*) Conditioned responding on the probe test in the identity unblocking paradigm. Exp: experimental group, receiving inhibition during reward outcome. ITI: control group, receiving inhibition during the intertrial interval. Asterisk indicates significant difference (*p* < 0.05). Data replotted from [1]. (*Right*) Model simulation of the value function.

**Figure 2:**
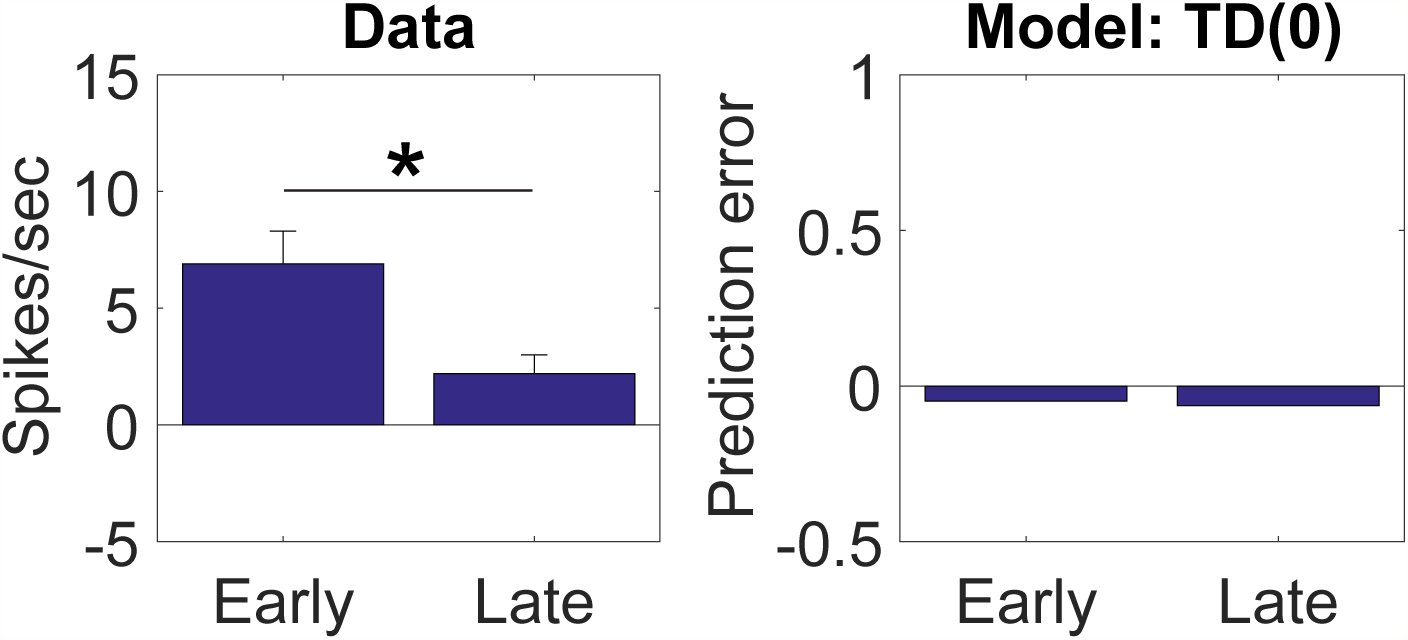
Dopamine neurons respond to changes in reward identity. (*Left*) Firing rate of dopamine neurons on trials that occurred early (first 5 trials) or late (last 5 trials) during an identity shift block. Data replotted from [2]. (*Right*) Model simulation of TD error.

**Figure 3:**
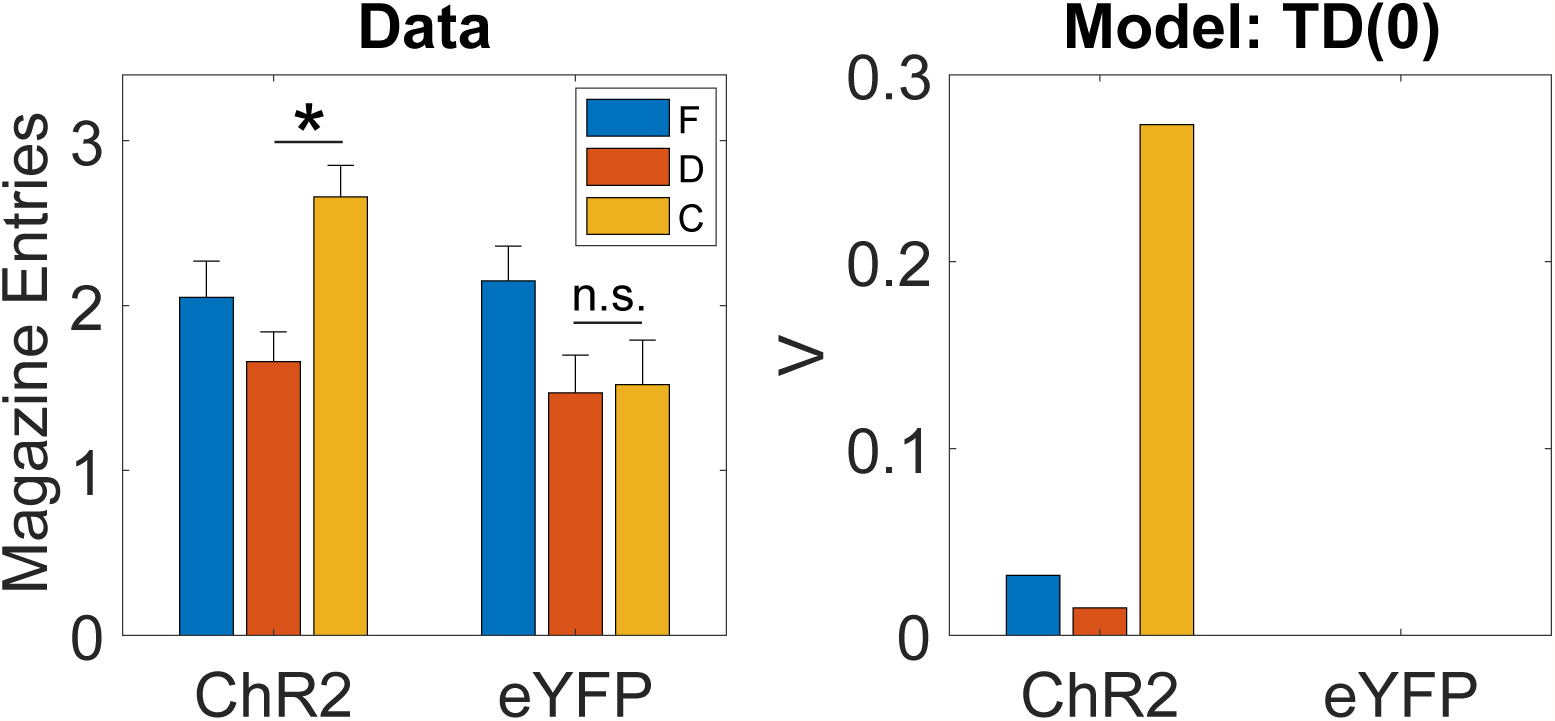
Dopamine transients are sufficient for learning stimulus-stimulus associations. (*Left*) Number of food cup entries occurring during the probe test for experimental (ChR2) and control (eYFP) groups in the sensory preconditioning paradigm. Data replotted from [3]. (*RIght*) Model simulation, using the value estimate as a proxy for conditioned responding. Note that V attached to the critical cue, C, is high in the simulation, much like the food cup responding in the probe test to this cue. This occurs because dopamine is paired with the cue, so it directly acquires a significant value. However, in this paradigm there is no direct link between C and the policy of going to the food cup. Thus, the success of TD(0) in this context in matching the empirical data is somewhat misleading.

**Figure 4:**
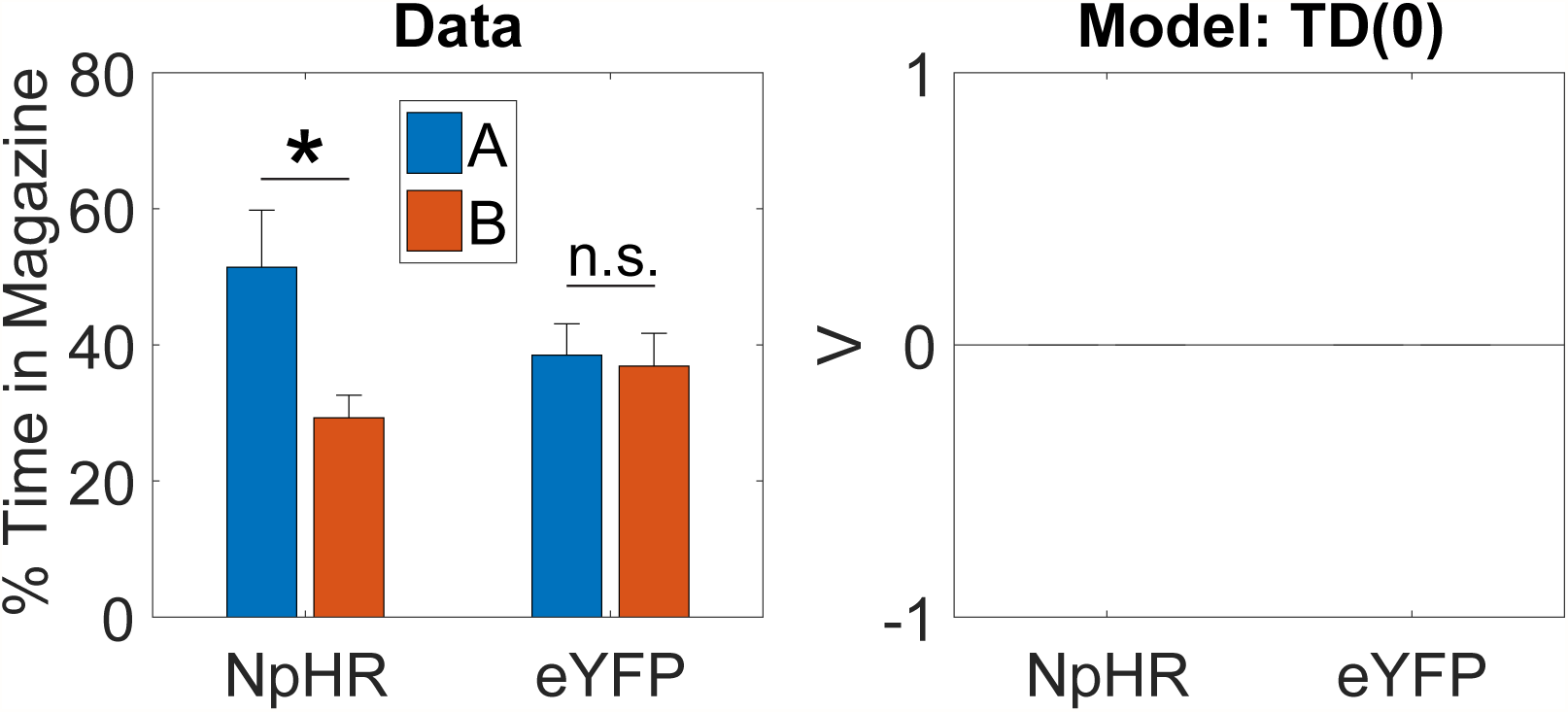
Dopamine transients are necessary for learning stimulus-stimulus associations. (*Left*) Number of food cup entries occurring during the probe test for experimental (NpHR) and control (eYFP) groups in the sensory preconditioning paradigm. Data replotted from [3]. (*Right*) Model simulation.

**Figure 5:**
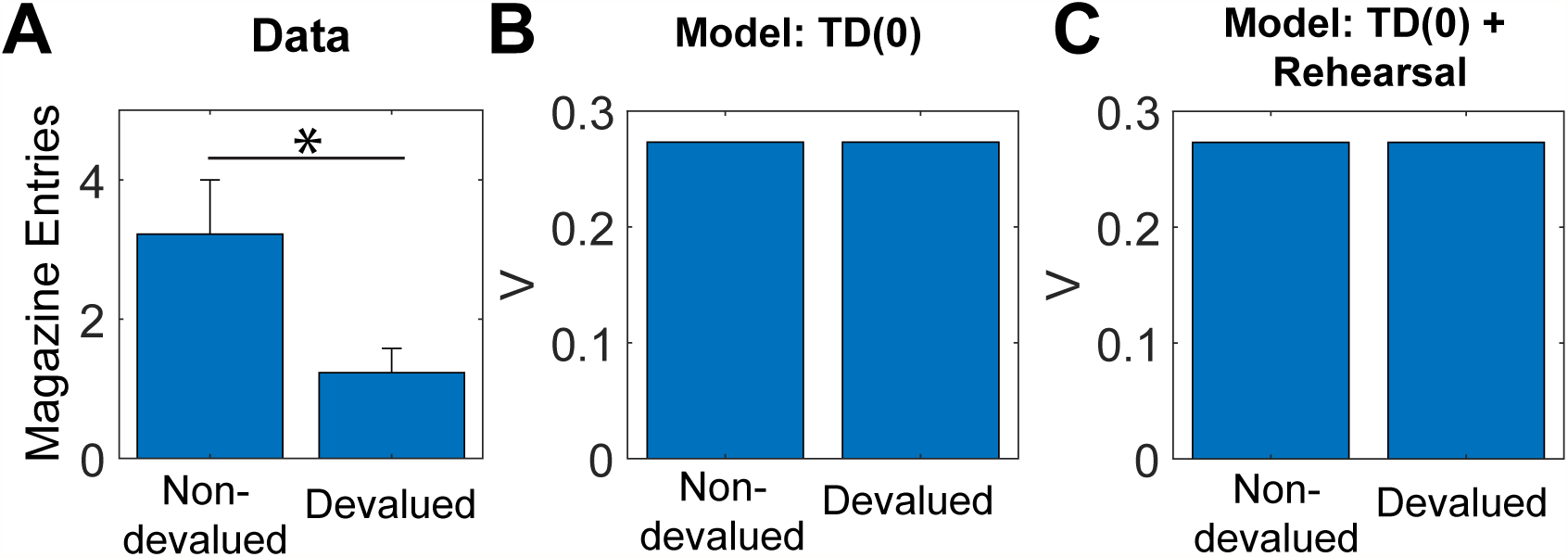
Behavior to preconditioned cue that is unblocked by activation of dopamine neurons is sensitive to devaluation of the predicted reward. Data (A, replotted from [3]) and model simulation (B) for conditioned responding to stimulus C in the probe test. Animals in the devalued group were injected with lithium chloride in conjunction with ingestion of the reward (sucrose pellets), causing a strong aversion to the reward. Animals in the nondevalued group were injected with lithium chloride approximately 6 hours after ingestion of the reward. (C) A version of the model with rehearsal of stimulus X during reward devaluation was able to capture the devaluation-sensitivity of animals.

